# Prevalence and socioeconomic determinants of development delay among children in Ceará, Brazil: a population-based study

**DOI:** 10.1101/597252

**Authors:** Luciano L. Correia, Hermano A. L. Rocha, Christopher R. Sudfeld, Sabrina G. M. O. Rocha, Álvaro J. M. Leite, Jocileide S. Campos, Anamaria C. e Silva

## Abstract

**Objective:** To assess the prevalence of child development delay and to identify socioeconomic determinants.

**Study Design:** We conducted a population-based cross-sectional study of children 2 to 72 months of age residing in the state of Ceará, Brazil. In total, 3200 households were randomly selected for participation in the study and had child development assessed with the Ages and Stages Questionnaire (ASQ) version 3. Development delay was defined as a score less than −2 standard deviations below the median of the Brazilian ASQ standard. We present population-level prevalence of delay in five development domains and assess socioeconomic determinants.

**Results:** A total of 3566 children completed the ASQ development assessment of which 9.2% (95% CI: 8.1-10.5) had at least one domain with development delay. The prevalence of delay increased with age in all domains and males were at higher risk for communication, gross motor and personal-social development delays as compared to females (p-values <0.05). We found robust associations of indicators of socioeconomic status with risk of development delay; increasing monthly income and higher social class were associated with reduced risk of delay across all domains (p-values <0.05). In addition, children in poor households that participated in conditional cash transfer (CCT) programs appeared to have reduced risk of delay as compared to children from households that were eligible, but did not participate, in CCT programs.

**Conclusions:** There is a relatively high population-level prevalence of development delay in at least one domain among children 0-6 years of age in Ceará, Brazil. Integrated child development, social support, and poverty reduction interventions may reduce the population-level prevalence of development delay in Ceará and similar settings.

## INTRODUCTION

The period from birth to 5 years of age is a critical window for development.(1) Globally, it is estimated that almost 250 million children under five years of age were at risk of failing to reach their full development potential in 2010.(2) Developmental deficits during early childhood may persist and lead to poor schooling achievement and result in reductions in lifetime earnings.(1, 3, 4) As a result, alleviating early life adversity may support attainment health and poverty reduction goal in low-and middle-income countries (LMIC).

There is rapidly growing literature on child development in LMIC settings; however, few studies have evaluated population representative child development and risk factors for delay. Population-level studies in America, Europe and Asia found the prevalence of development delay among children varies from 5 to 15%.(5-8) To our best knowledge, there are no population-based studies in Brazil. Population representative data is needed in Brazil to guide strategies to improve early childhood development to support meeting Sustainable Development Goal target 4.2 which call on countries to ensure all children have access to quality early childhood development, care and pre-primary education.(9)

We present a large cross-sectional and population-representative study of child development among children 0-6 years of age in the state of Ceará, Brazil. We determined the prevalence of children development delay as assessed by the Ages and Stages Questionnaire version 3 and assessed age, sex, and socioeconomic correlates of delay. Our study is intended to inform priority interventions and the need for integration of social protection and poverty-reduction programs to equitably improve child development in Ceará, Brazil.

## METHODS

### Study design and population

We analyzed cross-sectional data from the *Pesquisa de Saúde Materno Infantil do Ceará* (PESMIC) study. Full details of the PESMIC study methods can be found elsewhere.(10) Briefly, the PESMIC is a population-based study on maternal and child health of preschool children up to 72 months of age living in the state of Ceará, in northeastern Brazil. Population-based surveys were conducted in 1987, 1990, 1994, 2001, 2007 and 2017 using the same methods. For this article, we used data from 2017 which collected child development data.

Ceará is one of the poorest states in Brazil, with a population of 9 million inhabitants living in a semiarid climate. Fortaleza (2.3 million inhabitants) is the capital and urban commercial center of Ceará. The study area also includes rural areas of Ceará where subsistence farming is dominant. *Bolsa Família* is the world’s largest conditional cash transfer (CCT) program, including almost 14 million families or 55 million individuals in Brazil (more than a quarter of the entire population).(11) *Bolsa Família* is a direct income transfer program for families living in poverty (monthly income per capita between R$ 85.01, US$ 22.94, and R$ 170.00, US$ 45.87) and extreme poverty (monthly income per capita of up to R$ 85.00, US$ 22.94), created in 2003, combining several other fragmented social assistance programs. The average grant is about US$75 per month. In Ceará, 1,043,476 families are beneficiaries of the *Bolsa Família* program.

The PESMIC used cluster sampling based on the Brazilian Institute of Geography and Statistics (IBGE) census tracts with stratification between the state capital Fortaleza, and the rural areas. Census tracts were constructed from the division of each municipality into geographic areas with a stable population of 300 families. To ensure the study population was representative, municipalities, census tracts and households were randomly selected. Once a census tract was defined and its corresponding map obtained, 20 houses were randomly selected. The starting point of the cluster (the first home to be visited) was randomly selected utilizing ArcGIS^®^ software, GIS Inc. Households were visited consecutively, in a counterclockwise fashion. Shops and abandoned buildings were excluded and replaced; and in the case of absent families, up to three return visits were conducted in an attempt to obtain data.

Data used in the present study was collected from August to November of 2017, using three questionnaires covering household, maternal and child factors. The questionnaires were reviewed daily by field supervisors to identify and correct errors. During the fieldwork, data of a subsample (10% of children) were reassessed by supervisors for quality control. In the 2017 PESMIC the sample size was defined in 160 census tracts randomly selected. This sample comprised 3200 households. All children from 2 to 72 months old in households were eligible. In each household, information was obtained about all children through their mother or primary caregiver, and the children’s anthropometric measurements were taken.

### Child development assessment

To assess children’s development status, we used the Ages and Stages Questionnaire version 3,(12) which was validated in Brazil (ASQ-BR) (13) and has been used prior studies.(14) The ASQ consists of a series of questionnaires divided into 20 different age ranges; which seeks to evaluate children aged between 2 and 66 months (five and a half years). Five domains of child development weremeasured in the ASQ-BR subsections: communication, broad motor coordination, fine motor coordination, problem solving, and personal/social.(12) Interviewers were trained for 20 hours by medical professionals who were experienced with the ASQ-BR.

### Socioeconomic Factors

We collected data on multiple indicators of socioeconomic status. Head of households were asked to report their monthly income in *Reais* (Brazilian currency) and participation status in *Bolsa Família*. Social class was determined by the Brazilian Criteria “Critério Brasil”, which was developed using a representative sample of the family budget research (POF), carried out by the IBGE, classifying 55,970 domiciles based on household assets.(15) Food insecurity was assessed through the application of the United States Department of Agriculture (USDA) questionnaire modified by the experience of focal groups in Brazil.(16) The instrument consists of 15 central closed questions, with a yes /no response on the experience in the last three months of food insufficiency at its various levels of intensity, ranging from the apprehension that food may be lacking until the experience of passing all day without eating. Each affirmative answer of the questionnaire is equivalent to one point, varying the score from 0 to 15 points, considering the value zero as the safety condition; 1-5 points as mild insecurity; 6-10 points as moderate insecurity and 11-15 points as severe insecurity.

### Statistical Analysis

First, we tabulated total domain scores for each child using standard ASQ methods. For the final scores, if more than two items in an area were skipped, that particular domain was not scored. If one or two items in an area were skipped, we provided an adjusted score by calculating the average score for the completed items in that area, then assigning the skipped item(s) the average score.(12) An adjustment was made for children who were born premature and aged up to 24 months. We then standardized development scores for children greater than five months of age to the ASQ-BR standard which was created using 45,640 Brazilian children, stratified by age and sex.(17) We used American standardized score cut-offs for children from 2 to 4 months old.(18) As suggested in ASQ manual and in the literature, we considered < −2 standard deviation as a confirmatory delay and < −1 standard deviation as risk of delay.(12, 19) Calculated population-prevalence were adjusted for the sampling design that included stratification and cluster-based sampling. We used χ^2^tests adjusted for cluster-sampling to assess statistical significance of differences by socioeconomic indicators. Data were entered twice using EpiInfo 2000 and tested for concordance and analyzed using SPSS Version 23 (SPSS Statistics for Windows, Version 23.0. IBM Inc).

### Ethics

Written consent was obtained from participating women. Written consent for child participation was given by their mothers and consent for participating adolescent minors was obtained from parents or guardians. The survey was approved by the Research Ethics Committee in Brazil, under number 73516417.4.0000.5049.

### Data availability

Data made available to all interested researchers upon request.

## RESULTS

We contacted 3,200 households for participation in the study, of which all agreed to participate, resulting in a sample size of 3,566 children 0-6 years of age. Table 1 presents the demographic and socioeconomic characteristics of the study population. A total of 1,594 (44.6%) children resided in urban areas of Ceará. The mean monthly family income was1090.4 ±1017.9 *Reais* (about US$ 280) and 78% of the sample was in the lowest Brazilian socioeconomic class. In addition, 58% of the children resided in households that were food insecure and 61% of the families participated in conditional cash transfer programs.

**TABLE 1.** Characteristics of households and participants who completed the Ages and Stages Questionnaire

| Characteristics |  | N (%) or mean ± standard deviation<sup>1</sup> |
| --- | --- | --- |
| Child Sex | Male | 1786 (50) |
|  | Female | 1779 (49.9) |
| Residence | Urban | 1594 (44.6) |
|  | Rural | 1972 (55.3) |
| Child age in months |  | 31.8 ± 23.1 |
| Monthly income in Reais |  | 1090.4 ± 1017.9 |
| Social Class | Lower | 2744 (78.4) |
|  | C2 | 539 (15.4) |
|  | C1 | 161 (4.6) |
|  | B2 | 51 (1.4) |
|  | B1 | 5 (0.1) |
|  | Higher | 0 (0) |
| Food insecurity |  | 1840 (58) |
| Head of Household education | Up to 4 years | 2426 (73.3) |
|  | 5 - 8 years | 690 (20.8) |
|  | More than 8 years | 191 (5.7) |
| Participation in CCT programs | Yes, participate | 1943 (54.4) |
|  | No, but is eligible | 224 (6.2) |
|  | Not eligible | 1399 (39.2) |

1 A total of 3,200 households participated including 3,566 children

The prevalence of development delay (ASQ-3 domain score < −2 SD) by domain, age and sex is presented in Table 2. Overall, the population-level prevalence of communication delay was 2.0% (95 CI: 1.6-2.7), gross motor 3.8% (95 CI: 3.2-4.6), fine motor coordination 2.7% (95% CI: 2.2-3.5), problem-solving 2.8% (95 CI: 2.3-3.5) and personal-social 2.5% (95 CI: 2.0-3.2). We found that the prevalence of delay for all domains was greater for children who were 36-72 months of age as compared to those who were less than 36 months of age (p-values <0.05). In addition, we determined that males were at higher risk for communication, gross motor and personal-social development delays as compared to females (p-values <0.05).

**Table 2.** Population prevalence of development delay by Ages and Stages Questionnaire domain, child age, and child sex in Ceará, Brazil

|  |  | Communication | Gross motor | Fine motor | Problem-solving | Personal-social |
| --- | --- | --- | --- | --- | --- | --- |
|  | Total |  |  |  |  |  |
|  | N | N% (95% CI) | N% (95% CI) | N% (95% CI) | N% (95% CI) | N% (95% CI) |
| All children 2 to 66 months of age |  |  |  |  |  |  |
| All | 3284 | 2.0% (1.6-2.7) | 3.8% (3.2-4.6) | 2.7% (2.2-3.5) | 2.8% (2.3-3.5) | 2.5% (2.0-3.2) |
| Male | 1652 | 2.6% (1.9-3.5)* | 4.8% (3.9-6.0)* | 3.1% (2.4-4.1) | 3.1% (2.4-4.1) | 3.2% (2.4-4.2)* |
| Female | 1632 | 1.5% (0.9-2.3)* | 2.8% (2.0-3.7)* | 2.4% (1.7-3.4) | 2.5% (1.8-3.4) | 1.8% (1.2-2.8)* |
| 0 to 24 months |  |  |  |  |  |  |
| All | 1413 | 1.3% (0.8-2.1) | 3.4% (2.6-4.6) | 1.1% (0.7-1.3) | 1.7% (1.2-2.6) | 0.8% (0.5-1.4) |
| Male | 710 | 1.2% (0.6-2.3) | 3.3% (2.1-5) | 0.9% (0.4-1.3) | 2.4% (1.5-3.9)* | 0.3% (0.1-1.2)* |
| Female | 713 | 1.3% (0.6-3) | 3.6% (2.5-5.2) | 1.3% (0.7-2.4) | 1% (0.5-2.1)* | 1.3% (0.7-2.5)* |
| 24 to 36 months |  |  |  |  |  |  |
| All | 645 | 2.4% (1.4-3.9) | 2.5% (1.4-4.4) | 0.5% (0.2-1.5) | 1.4% (0.8-2.7) | 2.7% (1.7-4.3) |
| Male | 331 | 3.7% (2.1-6.5)* | 2.2% (1-4.9) | 1% (0.3-2.8) | 1.6% (0.7-3.6) | 4.3% (2.5-7.2)* |
| Female | 314 | 1.0% (0.4-2.9)* | 2.9% (1.5-5.7) | 0.0% (0-0) | 1.3% (0.5-3.4) | 1.0% (0.4-3)* |
| 36 to 72 months |  |  |  |  |  |  |
| All | 1226 | 2.9% (2.2-3.9) | 5% (3.9-6.4) | 5.9% (4.6-7.6) | 4.9% (3.8-6.5) | 4.5% (3.5-6) |
| Male | 611 | 3.8% (2.6-5.5) | 8.2% (6.3-10.7)* | 6.9% (5.1-9.3) | 5.0% (3.5-7) | 6.1% (4.4-8.5)* |
| Female | 615 | 2.0% (1.2-3.4) | 1.8% (1.1-3.2)* | 4.9% (3.4-7.2) | 4.9% (3.4-7.1) | 3% (1.9-4.7)* |

\* Statically significant difference between males and females (p value lower than 0.05).
¶All were statistically significant difference in combined group by age (0-36 and 36-72 months)

We then present the cumulative number of domains per child with developmental delay (< −2 SD domain score) and also for risk of developmental delay (<-1 SD domain score) in Table 3. In total, 9.2% (95%: 8.1-10.5) of all children sampled had at least one domain with development delay. The prevalence of at least one domain with development delay was significantly greater among males (11.0%; 95% CI: 9.5-12.6%) as compared to females (7.5%; 95% CI: 6.1-9.1) (p-value = 0.02). The overall prevalence of risk of developmental delay (<-1 SD) in at least one domain was 24.3% (95% CI: 22.5-26.2) with a significantly higher prevalence among males (28.1%; 95% CI: 25.8-30.5) as compared to females (20.5%; 95% CI: 18.3-22.8) (p-value <0.001).

**Table 3.**
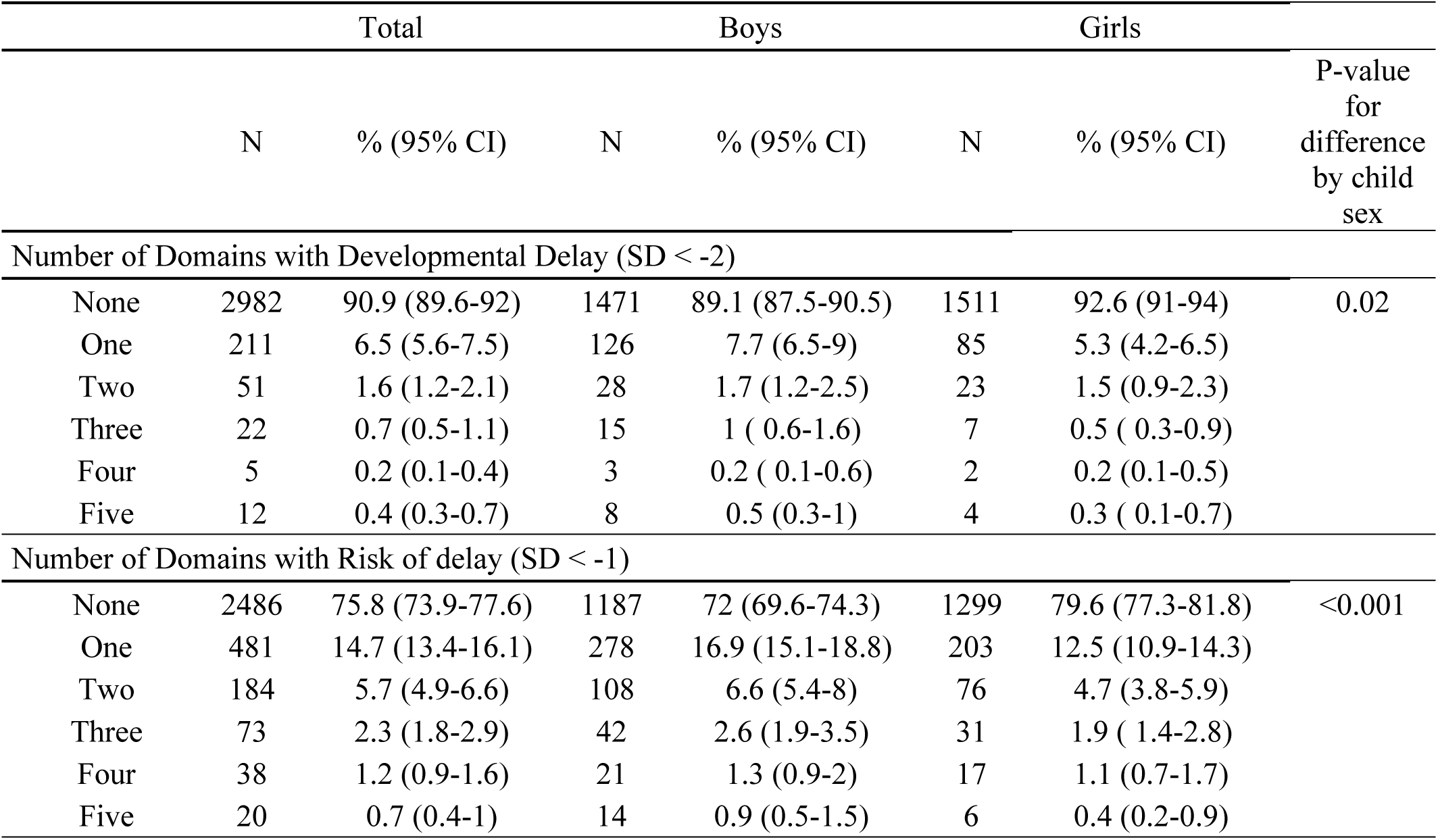
Cumulative number of domains with developmental delay per child out of the five Ages and Stages Questionnaire domains (communication, gross motor, fine motor, problem-solving, and personal-social domains) overall and stratified by child sex.

We present the prevalence of development delay by socioeconomic characteristics in Table 4. We found that increasing monthly income quintile was associated with reduced risk of delay in the communication, gross motor and fine motor domains and for delay in any domain (p-values <0.05). In addition, social class was associated with gross motor and fine motor delays (p-values <0.05), while food insecurity was associated with communication delay (p-value = 0.02). Among individuals eligible for conditional cash transfers, it appeared those who participated had lower risk of development delay in communication, gross motor and personal-social domains; however, the results did not reach statistical significance (p-values >0.05). There was no association of the head of household education with development delay in any domain. Similar results were found for risk of development delay, presented in table 5.

**Table 4.**
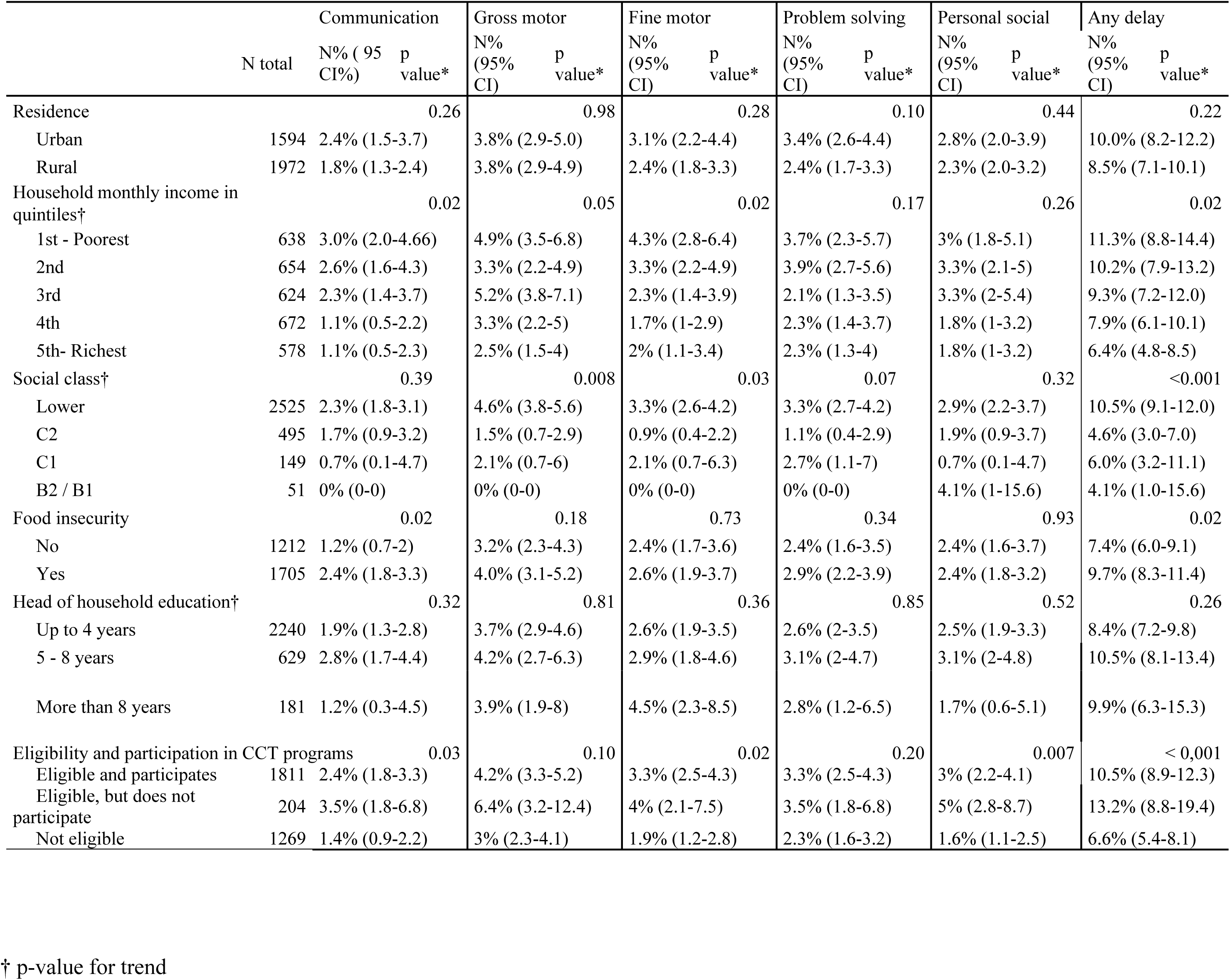
Prevalence of child development delay by domain stratified by socioeconomic factors.

**Table 5.**
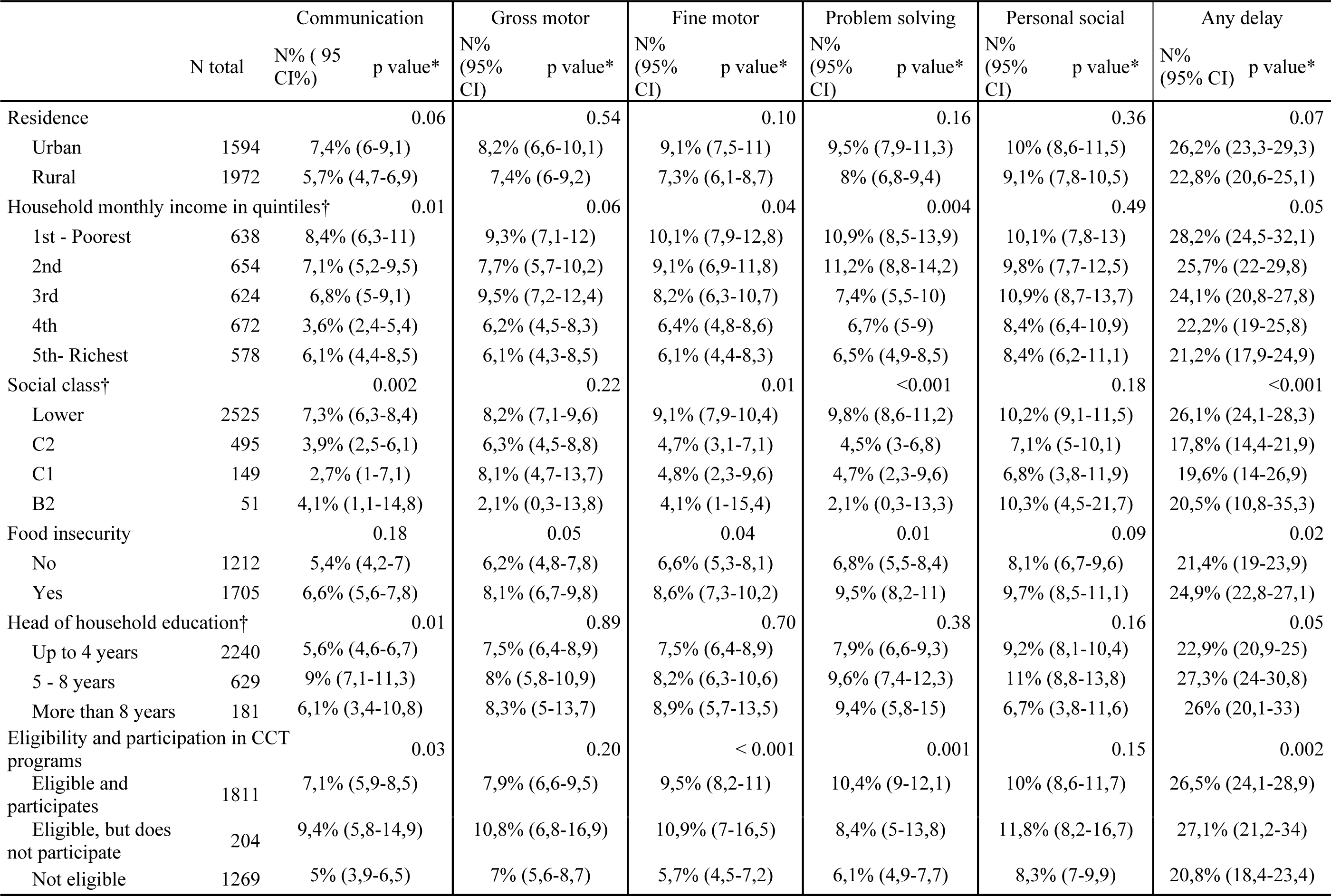
Prevalence of risk of child development delay by domain stratified by socioeconomic factors.

## DISCUSSION

We conducted a population-based study of development delay among children 2 months to 6 years of age in Ceará, Brazil. We found that 2-3% prevalence of development delay within each of the five domains and a 9.2% prevalence of development delay in at least one of the five domains assessed. Within each domain, the percent of children with delay was comparable to the Brazilian reference population as the percent of children < −2 SD is expected to be 2.5%. Nevertheless, we found that within Ceará, older children were at increased risk of delay across domains and male children had greater risk of delay in the gross motor, problem-solving, and personal-social domains. We also found that lower socioeconomic status, as assessed by monthly income and social class, was strongly associated with increased risk of development delay. In addition, there was some indication of lower prevalence of delay in poor families that participated in CCT programs as compared to those who were eligible but did not participate.

The prevalence of development delay in any of the five development domains in our study (9.2%) which is in line with existing literature in other settings.(5, 6, 8) In non-population representative samples in Brazil, the prevalence of development delay was reported to range from 21.4% to 46.3%, using different appraisal tools and in different ages.(20, 21) To the best of our knowledge this is the first comprehensive population-based study of the risk of child development delay in Brazil. Brazil has a development monitoring program that is estimated to achieve only 4.6% detection rates of cases and referral in some states.(20) Thus, the relatively high population prevalence of any delay and risk of delay and the low reach of monitoring programs suggests the need for evaluating the implementation of current child development policies in Ceará.

We also identified that male children in Ceará had a higher prevalence of delay in the motor, problem-solving, and personal-social domains as compared to females. This finding is in line with other studies that identified higher risk of suboptimal development, particularly in verbal and reading abilities, for males as compared to females.(5, 22-27) We are not able to determine mechanisms leading to the sex-differences but we hypothesize differences in parental stimulation by sex and age in Brazilian culture may be a contributor(28). In our study, the magnitude of the difference between males and females appeared to increase after 36 months of age. By 3 years of age, children follow parental instructions and may partake in different play activities that can lead to sex differences(26). In some studies, greater access to healthcare and child development services has been proposed to explain the higher prevalence of child development delay among males; however, assessment with the age and sex standardized ASQ-BR in this study does not support this mechanism. As a result, child development interventions that have some degree of tailoring by child sex may provide greater impact in this setting.

In this study we also found that social class, monthly household income and food security were robustly associated with development delays. There is a large literature on the link between poverty and child development.(29-32) Poverty may impact the child development in through a multitude of direct and indirect pathways. Poverty can directly lead to family stress and parental deprivation that leads to suboptimal development outcomes.(33) The chronification of poverty is more strongly associated with poor development than the acute poverty exposure, due to the sustained effect of this stress(34). In support, in our study we found that social class, had a stronger association with development and with risk of delay as compared to monthly income in some domains.(35, 36) This is particularly important because the home environment is more closely linked to the cognitive development of young preschool children than to that of school-age children.(30) In addition, poorest children often have less access to schools and to stimulating environments.(37) In Brazil two cohorts identified an association with family income and child development.(21) As a result, there is convincing evidence that targeting low SES households will be essential to improve population-level child development and reaching SDG target 4.2.

Consistent with the SES findings, we found that children who resided in households that were not eligible for CT programs had lower risk of delay than those who lived in eligible households. Nevertheless, we also found that among CT eligible households, children whose family participated in the programs tended to have lower risk of delay as compared to those whose family did not participate. This suggests that there may be some effect of the CT programs on child development outcomes; however, our observational analyses cannot be considered causal. The evidence on the effect of conditional and unconditional cash transfers on child development and general health outcomes is mixed(37, 38). In a study of *Oportunidades* program in Mexico, participating in the conditional cash transfer programs was associated with better cognition and language development and this finding was confirmed in in Nicaragua and Uganda.(32, 36, 39, 40) In Ceará, the *Bolsa Família* program’s conditionalities include attendance to antenatal care, children under 7 years of age to receive all recommended vaccines and all children attend routine health and growth monitoring visits.(41) Conditional cash transfers like *Bolsa Familia* may create positive externalities for improved child development among low SES households; however, additional research is needed to document if the existing CT programs provide child development benefits in Brazil.

This study has some limitations. First, the observational and cross-sectional design of the study does not afford analysis of child development trajectories over time and does not allow for direct determination of causal relationships. In addition, we used the ASQ-3, which is a validated screening tool, but is not diagnostic for child development delay. Further, while the study was designed to be population representative of children in the State of Ceará, the prevalence of delay and determinants are not likely generalizable to all Brazilian children.

Overall, we found that ∼10% of children under 6 years of age in Ceará, Brazil had delay in at least one development domain. In addition, the prevalence of delay increased with age which supports the need for early intervention. Males were also at higher risk for delay than females which may have some programmatic and relevance to design of stimulation and play interventions. We also found some evidence that cash transfer programs may ameliorate some of the risk of development delay among children in poor households. As a result, integrated child development interventions that also address underlying poverty-related risk factors and inequity may significantly reduce the population prevalence of development delay in Ceará and similar settings. Implementation research is needed to determine how to best deliver comprehensive child development interventions that are integrated with social protection and poverty reduction interventions.

## Conflicts of Interest and Source of Funding

All authors have no conflicts of interest to declare. The study was supported by the Fundação Cearense de Apoio ao Desenvolvimento Científico e Tecnológico (<PPSUS CE-FUNCAP/SESA/MS/CNPq 13506703-0>) and the Edital Jovens Doutores for postdoctoral support of HAL Rocha.

## Data sharing statement

Data will be made available upon request and further analysis.

There are no prior publications or submissions with any overlapping information, including studies and patients.

## Acknowledgments

To all participants of the study and specially to all mothers that sometimes even under unfavorable environmental, emotional and/or social conditions agreed to tell us their story.

## LIST OF ABBREVIATIONS

PESMIC: Pesquisa de Saúde Materno Infantil do Ceará
IBGE: Brazilian Institute of Geography and Statistics
USDA: United States Department of Agriculture
ANC: Antenatal care
LMIC: low- and middle-income countries
SES: socioeconomic status
SDG: Sustainable development goals
CTP: Cash Transfer Program

## REFERENCES

1. Grantham-McGregor S, Cheung YB, Cueto S, Glewwe P, Richter L, Strupp B. Developmental potential in the first 5 years for children in developing countries. The Lancet. 2007;369(9555):60–70.

2. Lu C, Black MM, Richter LM. Risk of poor development in young children in low-income and middle-income countries: an estimation and analysis at the global, regional, and country level. The Lancet Global health. 2016;4(12):e916–e22.

3. Smith JP. The impact of childhood health on adult labor market outcomes. The review of economics and statistics. 2009;91(3):478–89.

4. Fink G, Peet E, Danaei G, Andrews K, McCoy DC, Sudfeld CR, et al. Schooling and wage income losses due to early-childhood growth faltering in developing countries: national, regional, and global estimates, 2. The American journal of clinical nutrition. 2016;104(1):104–12.

5. Valla L, Wentzel-Larsen T, Hofoss D, Slinning K. Prevalence of suspected developmental delays in early infancy: results from a regional population-based longitudinal study. BMC Pediatrics. 2015;15(1):215.

6. Demirci A, Kartal M. The prevalence of developmental delay among children aged 3– 60 months in Izmir, Turkey. Child: Care, Health and Development. 2016;42(2):213–9.

7. Sajedi F, Vameghi R, Kraskian Mujembari A. Prevalence of undetected developmental delays in Iranian children. Child: Care, Health and Development. 2014;40(3):379–88.

8. King TM, Rosenberg LA, Fuddy L, Mcfarlane E, Sia C, Duggan AK. Prevalence and Early Identification of Language Delays Among At-Risk Three Year Olds. Journal of Developmental & Behavioral Pediatrics. 2005;26(4):293–303.

9. Griggs D, Stafford-Smith M, Gaffney O, Rockström J, Öhman MC, Shyamsundar P, et al. Sustainable development goals for people and planet. Nature. 2013;495:305.

10. Correia LL, Rocha HAL, Rocha SGMO, Nascimento LSd, Silva ACe, Campos JS, et al. Methodology of Maternal and Child Health Populational Surveys: A Statewide Cross-sectional Time Series Carried Out in Ceará, Brazil, from 1987 to 2017, with Pooled Data Analysis for Child Stunting. Annals of Global Health. 2019;85(1).

11. Pathways to Citizen Accountability: Brazil’s Bolsa Família AU - Sugiyama, Natasha Borges. The Journal of Development Studies. 2016;52(8):1192–206.

12. Squires J, Bricker DD, Twombly E. Ages & stages questionnaires: Paul H. Brookes Baltimore, MD; 2009.

13. Filgueiras A, Landeira-Fernandez J. Adaptação transcultural e avaliação psicométrica do Ages and Stages Questionnaires (ASQ) em creches públicas da cidade do Rio de Janeiro. Psicologia, PUC-Rio Rio de Janeiro: PUC-Rio. 2011.

14. Fioravanti-Bastos ACM, Filgueiras A, Moura MLSd. Evaluation of the Ages and Stages Questionnaire-Brazil by Early Childhood professionals. Estudos de Psicologia (Campinas). 2016;33:293–301.

15. Kamakura W, Mazzon J. Critérios de estratificação e comparação de classificadores socioeconômicos no brasil 2016. 55–70 p.

16. Corrêa A, Escarilha R, Sampaio M, Panigassi G, Maranha L, Bergamasio S. Acompanhamento e avaliação da segurança alimentar de famílias brasileiras: validação de metodologia e de instrumento de coleta de informação. Urbano/rural Relatório técnico: versão preliminar Brasília (DF): Ministério da Saúde: Organização Pan-Americana da Saúde. 2004.

17. Filgueiras A, Pires P, Maissonette S, Landeira-Fernandez J. Psychometric properties of the Brazilian-adapted version of the Ages and Stages Questionnaire in public child daycare centers. Early human development. 2013;89(8):561–76.

18. Janson H, Squires J. Parent-completed developmental screening in a Norwegian population sample: a comparison with US normative data. Acta paediatrica. 2004;93(11):1525–9.

19. Veldhuizen S, Clinton J, Rodriguez C, Wade TJ, Cairney J. Concurrent validity of the Ages and Stages Questionnaires and Bayley Developmental Scales in a general population sample. Academic pediatrics. 2015;15(2):231–7.

20. Caminha MdFC, Silva SLd, Lima MdC, Azevedo PTÁCCd, Figueira MCdS, Batista Filho M. Vigilância do desenvolvimento infantil: análise da situação brasileira. Revista Paulista de Pediatria. 2017;35:102–9.

21. Halpern R, Barros AJ, Matijasevich A, Santos IS, Victora CG, Barros FC. Developmental status at age 12 months according to birth weight and family income: a comparison of two Brazilian birth cohorts. Cad Saude Publica. 2008;24 Suppl 3:S444–50.

22. Kramer JH, Delis DC, Kaplan E, O’Donnell L, Prifitera A. Developmental sex differences in verbal learning. Neuropsychology. 1997;11(4):577–84.

23. Berglund E, Eriksson M, Westerlund M. Communicative skills in relation to gender, birth order, childcare and socioeconomic status in 18-month-old children. Scandinavian journal of psychology. 2005;46(6):485–91.

24. Hyde JS, Linn MC. Gender differences in verbal ability: A meta-analysis. Psychological bulletin. 1988;104(1):53.

25. Westerlund M, Lagerberg D. Expressive vocabulary in 18-month-old children in relation to demographic factors, mother and child characteristics, communication style and shared reading. Child: Care, Health & Development. 2008;34(2):257–66.

26. Fenson L, Dale PS, Reznick JS, Bates E, Thal DJ, Pethick SJ, et al. Variability in Early Communicative Development. Monographs of the Society for Research in Child Development. 1994;59(5):i–185.

27. Flamant C, Müller J-B, Rozé J-C, Olivier M, Rouger V, Hanf M, et al. Relative contributions of prenatal complications, perinatal characteristics, neonatal morbidities and socio-economic conditions of preterm infants on the occurrence of developmental disorders up to 7 years of age. International Journal of Epidemiology. 2018;48(1):71–82.

28. Spessato BC, Gabbard C, Valentini N, Rudisill M. Gender differences in Brazilian children’s fundamental movement skill performance. Early Child Development and Care. 2013;183(7):916–23.

29. and JLA, Bennett NG, Conley DC, Li J. The Effects of Poverty on Child Health and Development. Annual Review of Public Health. 1997;18(1):463–83.

30. and RHB, Corwyn RF. Socioeconomic Status and Child Development. Annual Review of Psychology. 2002;53(1):371–99.

31. Blau DM. The effect of income on child development. Review of Economics and Statistics. 1999;81(2):261–76.

32. Brooks-Gunn J, Duncan GJ. The effects of poverty on children. The future of children. 1997:55–71.

33. McLoyd VC, Wilson L. Maternal behavior, social support, and economic conditions as predictors of distress in children. New Directions for Child and Adolescent Development. 1990;1990(46):49–69.

34. Duncan GJ, Brooks-Gunn J. Family poverty, welfare reform, and child development. Child development. 2000;71(1):188–96.

35. J. DG, Jeanne B-G. Family Poverty, Welfare Reform, and Child Development. Child Development. 2000;71(1):188–96.

36. J. DG, Jeanne B-G, Kato KP. Economic Deprivation and Early Childhood Development. Child Development. 1994;65(2):296–318.

37. de Walque D, Fernald L, Gertler P, Hidrobo M. Cash Transfers and Child and Adolescent Development. Child and Adolescent Health and Development 3rd edition: The International Bank for Reconstruction and Development/The World Bank; 2017.

38. Pega F, Liu SY, Walter S, Pabayo R, Saith R, Lhachimi SK. Unconditional cash transfers for reducing poverty and vulnerabilities: effect on use of health services and health outcomes in low- and middle-income countries. Cochrane Database of Systematic Reviews. 2017(11).

39. Fernald LCH, Gertler PJ, Neufeld LM. Role of cash in conditional cash transfer programmes for child health, growth, and development: an analysis of Mexico’s Oportunidades. The Lancet. 2008;371(9615):828–37.

40. Macours KS, Norbert Vakis, Renos. Cash Transfers, Behavioral Changes, And Cognitive Development In Early Childhood: Evidence From A Randomized Experiment.

41. Social. SEdD. Bolsa família > o que é > acesso a educação e saúde 2019 [Available from: http://mds.gov.br/assuntos/bolsa-familia/o-que-e/acesso-a-educacao-e-saude.

